# Spontaneous behaviors drive multidimensional, brain-wide activity

**DOI:** 10.1101/306019

**Authors:** Carsen Stringer, Marius Pachitariu, Nicholas Steinmetz, Charu Bai Reddy, Matteo Carandini, Kenneth D. Harris

## Abstract

Cortical responses to sensory stimuli are highly variable, and sensory cortex exhibits intricate spontaneous activity even without external sensory input. Cortical variability and spontaneous activity have been variously proposed to represent random noise, recall of prior experience, or encoding of ongoing behavioral and cognitive variables. Here, by recording over 10,000 neurons in mouse visual cortex, we show that spontaneous activity reliably encodes a high-dimensional latent state, which is partially related to the mouse’s ongoing behavior and is represented not just in visual cortex but across the forebrain. Sensory inputs do not interrupt this ongoing signal, but add onto it a representation of visual stimuli in orthogonal dimensions. Thus, visual cortical population activity, despite its apparently noisy structure, reliably encodes an orthogonal fusion of sensory and multidimensional behavioral information.

---

In the absence of sensory inputs, the brain produces structured patterns of activity, which can be as large or larger than sensory driven activity [1]. Ongoing activity exists even in primary sensory cortices, and have been hypothesized to reflect recapitulation of previous sensory experiences, or expectations of possible sensory events. This hypothesis is supported by studies that found similarities between sensory-driven and spontaneous firing events [2–4]. An alternative possibility is that ongoing activity could be related to behavioral and cognitive states. The firing of sensory cortical and sensory thalamic neurons correlates with behavioral variables such as locomotion, pupil diameter, and whisking [5–19]. Continued encoding of nonvisual variables when visual stimuli are present could at least in part explain the trial-to-trial variability in responses to repeated presentation of identical sensory stimuli [20].

The influence of trial-to-trial variability on stimulus encoding depends on its population-level structure. Variability that is independent between cells – such as the Poisson-like variability produced in balanced recurrent networks [21] – presents little impediment to information coding, as reliable signals can still be extracted as weighted sums over a large enough population. In contrast, correlated variability has consequences that depend on the form of the correlations. If correlated variability mimics the differences in responses to different stimuli, it can compromise stimulus encoding [22]. Conversely, variability in dimensions orthogonal to those encoding stimuli has no adverse impact on coding [23], and might instead reflect encoding of signals other than visual inputs.

## Spontaneous cortical activity reliably encodes a high-dimensional latent signal

To distinguish between these possibilities, we characterized the structure of neural activity and sensory variability in mouse visual cortex. We simultaneously recorded from 11,262 ± 2,282 (mean ± s.d.) excitatory neurons, over nine sessions in six mice using 2-photon imaging of GCaMP6s in an 11-plane configuration [24] (Figure 1A,B, Movie S1). These neurons’ responses to classical grating stimuli revealed robust orientation tuning as expected in visual cortex (Figure S1).

**Figure 1.**
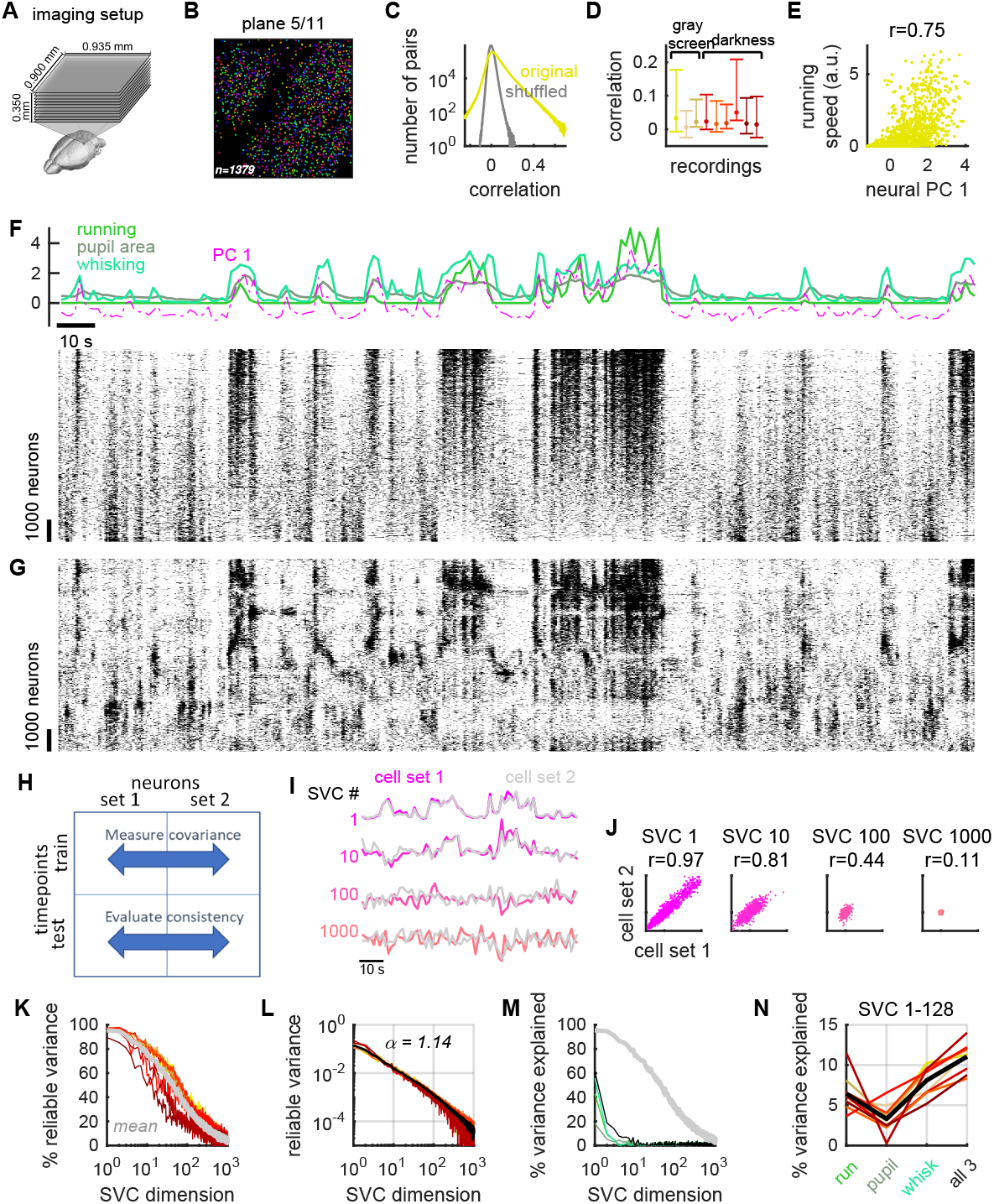
Structured ongoing population activity in visual cortex. (A) Two-photon calcium imaging of ~10,000 neurons in visual cortex using multi-plane resonance scanning of 11 planes spaced 35 *μ*m apart. (B) Randomly-pseudocolored cells detected in an example imaging plane. (C) Distribution of pairwise correlations in ongoing activity, computed in 1.2 second time bins (yellow). Gray: distribution of correlations after randomly time-shifting each cell’s activity. (D) Bars showing distribution of pairwise correlation coefficients for each recording (dot: mean; bar, range between 5th and 95th percentile). (E) First principal component (PC) of population activity, plotted versus running speed. Each point represents a 1.2 s time bin. (F) Top: example timecourse of running speed (green), pupil area (gray), whisking (light green), first PC (magenta dashed). Bottom: raster representation of ongoing population activity, with neurons sorted vertically by 1st PC weighting. (G) Same neurons as in (F), sorted by a manifold embedding algorithm to reveal multi-dimensional structure of population activity. (H) Shared Variance Component Analysis method for estimating the reliable variance spectrum of spontaneous neural activity. Dimensions of maximum covariance between two spatially-separated cell sets are estimated from half the recording (training timepoints), and their covariance on the test timepoints yields an asymptotically unbiased estimate of reliable variance. (I) Example time course of SVCs 1, 10, 100, and 1000 from each cell set in the test epoch. (J) Comparison of SVCs estimated independently from the two cell sets. Each dot represents a 1.2 s period. Pearson correlations (top) estimate fraction of that dimension’s variance reliably encoding latent signal. (K) Fraction of reliable variance for successive dimensions. Colors: different experiments; gray: grand mean. (L) Estimated reliable variance spectrum, showing a power law of exponent 1.14. (M) Percentage of each SVC’s total variance that can be reliably predicted from arousal variables. Shades of green: different arousal variables, color coded as in F. Light gray: percentage of reliable variance, as in K. (N) Percentage of total variance in first 128 dimensions explainable by arousal variables, individually or combined.

We began by analyzing spontaneous activity in mice free to run on an air-floating ball. Six of nine recordings were performed in darkness, but we did not observe differences between these recordings (shown in red on all plots) and recordings with gray screen (yellow on all plots). Mice spontaneously performed behaviors such as running, whisking, sniffing, and other facial movements, which we monitored with an infrared camera.

Ongoing population activity in visual cortex was highly structured (Figure 1C-F). Correlations between neuron pairs were reliable (Figure S2), and their spread was larger than would be expected by chance (Figure 1C,D), suggesting structured activity [25]. Fluctuations in the first principal component (PC) occurred over a timescale of many seconds (Figure S3), and were coupled to variations in arousal levels, as indicated by running, whisking, and pupil area. These arousal-related variables were strongly correlated with each other (Figure S4A,B), and together accounted for approximately 50% of the variance of the first neural PC (Figure 1E, Figure S4C). Correlation with the first PC was positive or negative in approximately similar numbers of neurons (57% ± 6.7% SE positive), indicating that two large sub-populations of neurons alternate their activity (Figure 1F). The slowness of these fluctuations implies a different underlying phenomenon to previously-studied “up and down phases” [3, 12, 26–28], which alternate at a much faster timescale (100-300 ms instead of 10-20 s) and correlate with most neurons positively. Indeed, up/down phases could not even have been detected in our recordings, which scanned the cortex every 400 ms.

Spontaneous activity had a high-dimensional structure, more complex than would be predicted by a single factor such as arousal (Figure 1G). This structure could be visualized by sorting the raster diagram so that nearby neurons showed strong correlations (see also Figure S5). Position on this continuum bore little relation to actual distances in the imaged tissue (Figure S6).

Despite its noisy appearance, spontaneous population activity reliably encoded a high-dimensional latent signal (Figure 1H-K). To show this, we devised a method to identify dimensions of neural variance that are reliably determined by common underlying signals, termed Shared Variance Component Analysis (SVCA). The method begins by dividing the recorded neurons into two spatially segregated sets, and dividing the recorded timepoints into two equal halves (training and test; Figure 1H). The training timepoints are used to find the dimensions in each cell set’s activity that maximally covary with each other. These dimensions are termed Shared Variance Components (SVCs). Activity in the test timepoints is then projected onto each SVC (Figure 1I), and the correlation between projections from the two cell sets (Figure 1J) provides an estimate of the reliable variance in that SVC (see Methods and Appendix). The fraction of reliable variance in the first SVC was 97% (Figure 1I,J), implying that only 3% of the variance along this dimension reflected independent noise. The reliable variance fraction of successive SVCs decreased slowly, with the 50th SVC showing approximately 50%, and the 512th showing 9% (Figure 1K). Thus, visual cortical spontaneous activity encodes a multidimensional latent signal, and appropriately weighted sums of 10,000 neurons suffice to accurately extract ~100 dimensions of it.

The magnitude of reliable spontaneous variance was distributed across dimensions according to a power law of exponent 1.14 (Figure 1L). This value is larger than the power law exponents close to 1.0 seen for stimulus responses [29], but still indicates a high-dimensional signal. The first 128 SVCs together accounted for 86% ± 1% SE of the complete population’s reliable variance, and 67% ± 3% SE of the total variance in these 128 dimensions was reliable. Arousal variables accounted for 11% ± 1% SE of the total variance in these 128 components (16% of their reliable variance), and primarily correlated with the top SVCs (Figure 1M,N).

## Ongoing neural activity encodes multidimensional behavioral information

Although arousal measures only accounted for a small fraction of the reliable variance of spontaneous population activity, it is possible that a larger fraction could be explained by higher-dimensional measures of ongoing behavior (Figure 2A-C, Movie S2). We extracted a 1,000-dimensional summary of the motor actions visible on the mouse’s face by applying principal component analysis to the spatial distribution of facial motion at each moment in time [30]. The first PC captured motion anywhere on the mouse’s face, and was strongly correlated with explicit arousal measures(Figure S4B), while higher PCs distinguished different types of facial motion. We predicted neuronal population activity from this behavioral signal using reduced rank regression: for any *N*, we found the *N* dimensions of the video signal predicting the largest fraction of the reliable spontaneous variance (Figure 2D).

**Figure 2.**
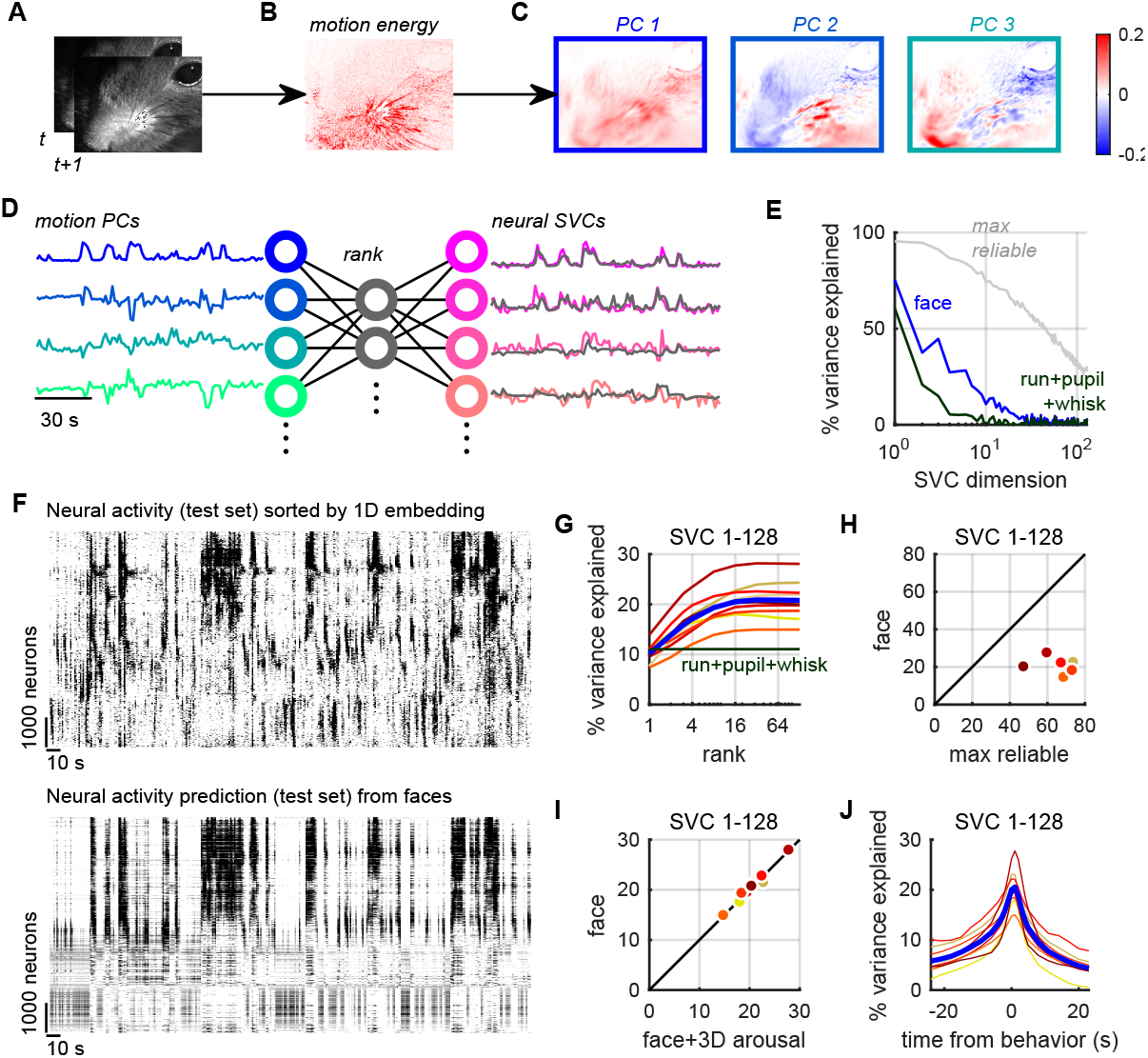
Multi-dimensional behavior predicts neural activity. (A) Frames from a video recording of a mouse’s face. (B) Motion energy, computed as the absolute value of the difference of consecutive frames. (C) Spatial masks corresponding to the top three principal components (PCs) of the motion energy movie. (D) Schematic of reduced rank regression technique used to predict neural activity from motion energy PCs. (E) Cross-validated fraction of successive neural SVCs predictable from face motion (blue), together with fraction of variance predictable from running, pupil and whisking (green), and fraction of reliable variance (the maximum explainable; gray; cf. Figure 1K). (F) Top: raster representation of ongoing neural activity in an example experiment, with neurons arranged vertically as in Figure 1G so correlated cells are close together. Bottom: prediction of this activity from facial videography (predicted using separate training timepoints). (G) Percentage of the first 128 SVCs’ total variance that can be predicted from facial information, as a function of number of facial dimensions used. (H) Prediction quality from multidimensional facial information, compared to the amount of reliable variance. (I) Adding explicit running, pupil and whisker information to facial features provides little improvement in neural prediction quality. (J) Prediction quality as a function of time lag used to predict neural activity from behavioral traces.

This multidimensional behavior measure predicted approximately twice as much variance as the simple arousal variables (Figure 2D-J, Movie S3). To visualize how multidimensional behavior predicts ongoing population activity, we compared a raster representation of raw activity (vertically sorted as in Figure 1G) to the prediction based on multidimensional videography (Figure 2F, see Figure S5 for all recordings). To quantify the quality of prediction, and the dimensionality of the behavioral signal encoded in V1, we focused on the first 128 SVCs (accounting for 86% of the population’s reliable variance). The best one-dimensional predictor extracted from the facial motion movie captured the same amount of variance as the best one-dimensional combination of whisking, running, and pupil (Figure 2G). However, prediction quality continued to increase with up to 16 dimensions of videographic information (and beyond, in some recordings), suggesting that visual cortex encodes at least 16 dimensions of motor information. These dimensions together accounted for 21%± 1% SE of the total population variance (31% ± 3% of the reliable variance; Figure 2H), substantially more than the three-dimensional model of neural activity using running, pupil area and whisking (11% ± 1% SE of the total variance, 17% ± 1% SE of the reliable variance). Moreover, adding these three explicit predictors to the video signal increased the explained variance by less than 1% (Figure 2I), even though the running signal was not obtained from the video camera. A neuron’s predictability from behavior was not related to its cortical location (Figure S7). The timescale with which neural activity could be predicted from facial behavior was ~1 s (Figure 2J), reflecting the slow nature of these behavioral fluctuations.

## Behaviorally-related activity is spread across the brain

The high-dimensional spontaneous activity patterns found in V1 were a reflection of activity patterns spread across the brain (Figure 3A-E). To show this, we performed large-scale electrophysiological recordings, using 8 Neuropixels probes [31] to simultaneously record from frontal, sensorimotor, visual and retrosplenial cortex, hippocampus, striatum, thalamus, and midbrain (Figure 3A,B). The mice were awake and free to rotate a wheel with their front paws. From recordings in three mice, we extracted 2,998, 2,768 and 1,958 units stable across ~1 hour of ongoing activity, and binned neural activity into 1.2 s bins, as for the imaging data.

**Figure 3.**
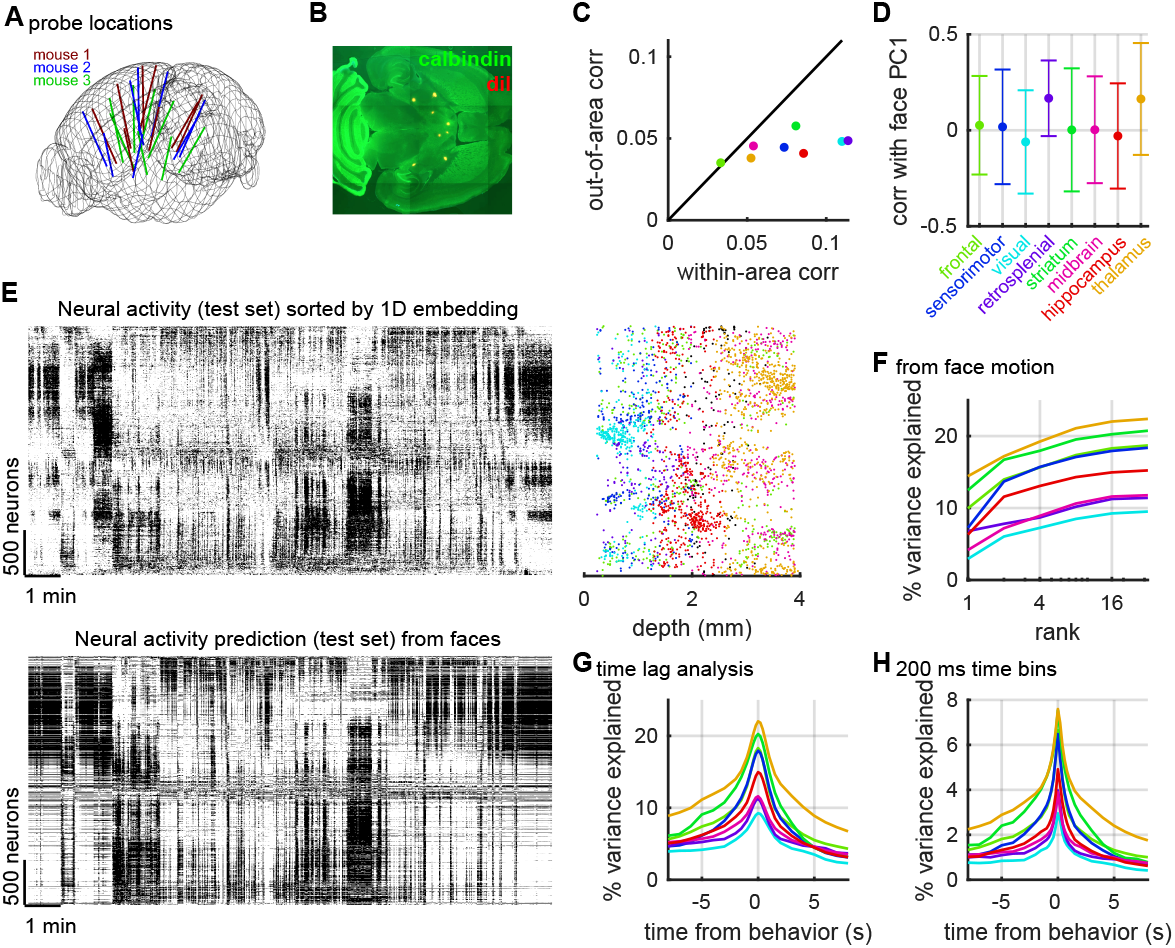
Behaviorally-related activity across the forebrain in simultaneous recordings with 8 Neuropixels probes. (A) Reconstructed probe locations of recordings in three mice. (B) Example histology slice showing orthogonal penetrations of 8 electrode tracks through a calbindin-counterstained horizontal section. (C) Comparison of mean correlation between cell pairs in a single area, with mean correlation between pairs with one cell in that area and the other elsewhere. Each dot represents the mean over all cell pairs from all recordings, color coded as in panel D. (D) Mean correlation of cells in each brain region with first principal component of facial motion. Error bars: standard deviation. (E) Top: Raster representation of ongoing population activity for an example experiment, sorted vertically so nearby neurons have correlated ongoing activity. Bottom: prediction of this activity from facial videography. Right: Anatomical location of neurons along this vertical continuum. Each point represents a cell, colored by brain area as in C,D, with x-axis showing the neuron’s depth from brain surface. (F) Percentage of population activity explainable from orofacial behaviors as a function of dimensions of reduced rank regression. Each curve shows average prediction quality for neurons in a particular brain area. (G) Explained variance as a function of time lag between neural activity and behavioral traces. Each curve shows the average for a particular brain area. (H) Same as G in 200ms bins.

Neurons were on average more strongly correlated with others in the same area, but substantial inter-area correlations also existed, suggesting non-localized patterns of neural activity (Figure 3C). All areas contained neurons positively and negatively correlated with the top facial motion PC, although thalamus and retrosplenial cortex contained predominantly arousal-preferring neurons (Figure 3D, p<10^−8^ two-sided Wilcoxon sign-rank test). Sorting the neurons by correlation again revealed a complex activity structure (Figure 3E). All brain areas contained a sampling of neurons from the entire continuum (Figure 3E, right), suggesting that a multidimensional structure of ongoing activity is distributed throughout the brain. This spontaneous activity spanned at least 128 dimensions, with 35% of the variance of individual neurons reliably predictable from population activity (Figure S8).

Similar to visual cortical activity, the activity of brainwide populations was partially predictable from facial videography (Figure 3F-H). Predictability of brain-wide activity again saturated around 16 behavioral dimensions, which predicted on average 16% of the total variance (42% of the estimated maximum possible) (Figure 3F). Other areas showed even stronger behavioral modulation than visual cortex, with neurons in thalamus predicted best (22% of total variance, 53% of estimated maximum). The timescale of videographic prediction was again broad: neural activity was best predicted from instantaneous behavior (17±24ms SE before behavior), but the predicted variance again decayed slowly over time lags of multiple seconds (Figure 3G). Taking advantage of the high temporal resolution of electrophysiological recordings, we reduced the analysis bin size from 1.2 seconds to 200 ms, revealing an additional sharp peak at short timescales across all areas (optimal timelag at 80±18 ms Figure 3H).

## Stimulus-evoked and ongoing activity overlap along one dimension

We next asked how ongoing activity and behavioral information relates to sensory responses (Figure 4A-B). For this analysis, we interspersed blocks of visual stimulation with flashed natural images (presented 1 per second on average) with extended periods of spontaneous activity (gray screen), while imaging visual cortical population activity (Figure 4A). During stimulus presentation, the mice continued to exhibit the same behaviors as in darkness, resulting in a similar distribution of facial motion components (Figure 4B).

**Figure 4.**
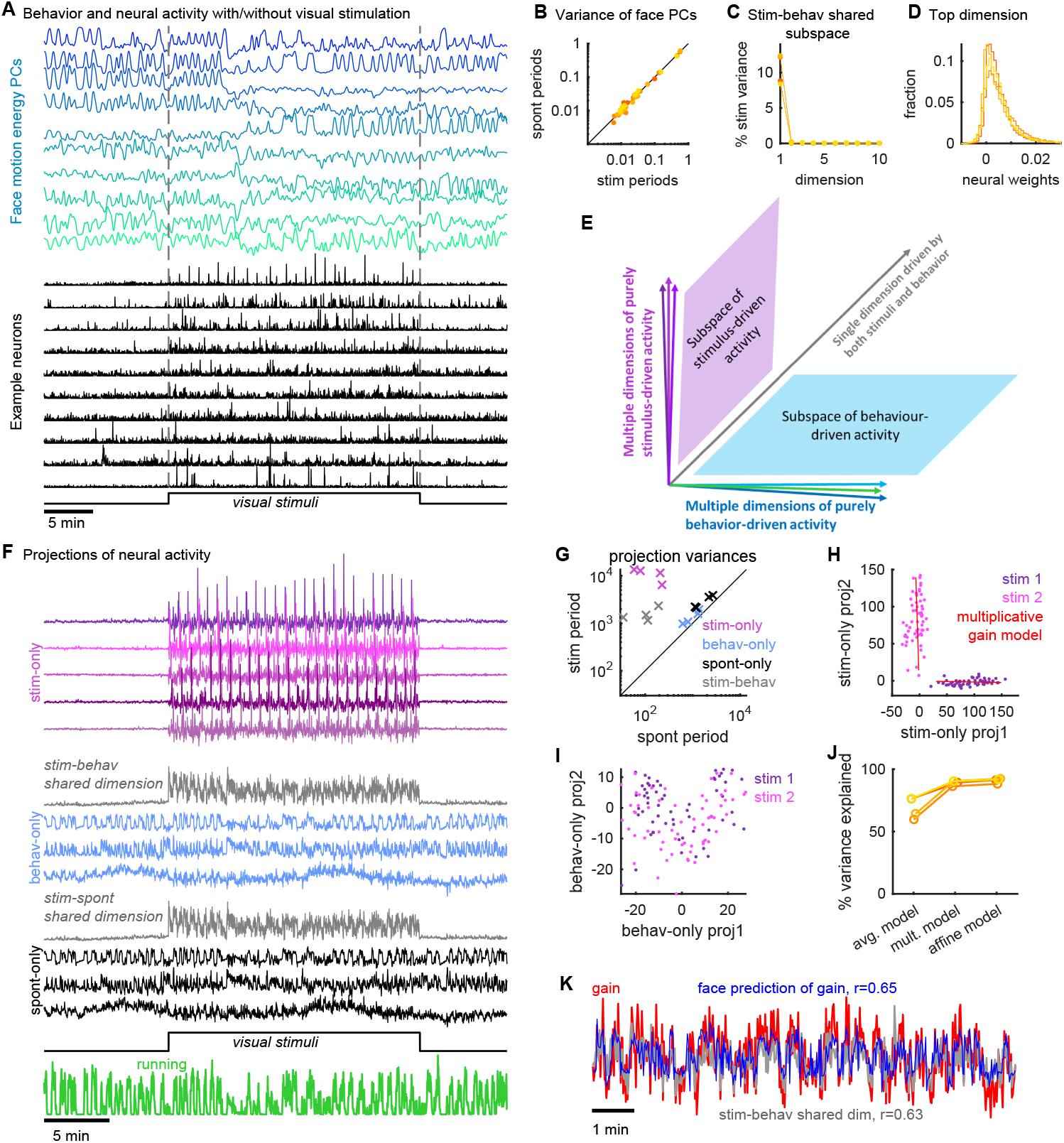
Neural subspaces encoding stimuli and spontaneous/behavioral variables overlap along one dimension. (A) Principal components of facial motion energy (top) and firing of ten example V1 neurons (bottom), before, during and after a period of visual stimulus presentation. (B) Comparison of facial motion energy with and without visual stimulation. Each point represents a single PC in a single experiment; color represents experiment identity. (C) Overlap of stimulus and behavioral subspaces. X-axis represents successive dimensions of overlap between the subspaces spanned by mean stimulus responses and facial prediction; y-axis represents fraction of stimulus-related variance in each dimension. (D) Distribution of cells’ weights on the single dimension of overlap between stimulus and behavior subspaces. Each curve represents the distribution of weights over all cells in an experiment. (E) Illustration of three sets of orthogonal dimensions in the subspace of firing patterns. Activity in multiple dimensions is driven by visual stimuli but not behavior (shades of magenta); multiple other dimensions are driven by behavior but not by stimuli (shades of cyan); a single dimension (gray; characterized in panels C,D) is driven by both. (F) Example of neural population activity projected onto these three sets of dimensions. Top: shades of magenta, projection onto stimulus-related dimensions. Middle: gray, projection onto single dimension related to both stimuli and behavior; blue, projection onto dimensions related to behavior alone. Bottom: similar analysis for all ongoing spontaneous dimensions, even if unrelated to facial behavior. (G) Amount of variance of each of the projections illustrated in F, during stimulus presentation and spontaneous periods. Each point represents summed variances of the dimensions in the subspace corresponding to the symbol color, for a single experiment. (H) Projection of population responses to repeats of two example stimuli into two dimensions of the stimulus-only subspace. Red lines: multiplicative gain model. (I) Similar plot for two dimensions of the behavior-only subspace. (J) Fraction of variance in the stimulus-only subspace explained by three models: constant response on each trial of the same stimulus (avg. model); multiplicative gain that varies across trials (mult. model); and a model with both multiplicative and additive terms (affine model). (K) The multiplicative gain on each trial can be predicted by the face motion PCs.

Representations of sensory and behavioral information were mixed together in the same cell population. There were not separate sets of neurons encoding stimuli and behavioral variables: the fraction of each neuron’s variance explained by stimuli and by behavior were only slightly negatively correlated (Figure S9; r = -0.18, p<0.01 Spearman’s rank correlation), and neurons with similar stimulus responses did not have more similar behavioral correlates (Figure S9; r = -0.005, p > 0.05).

Nevertheless, the subspaces encoding sensory and behavior information overlapped in only one dimension (Figure 4C-E). The space encoding behavioral variables contained 11% of the total stimulus-related variance, 96% of which was contained in a single dimension (Figure 4C) with largely positive weights onto all neurons (85% positive weights, Figure 4D). Similarly, the space of ongoing activity, defined by the top 128 principal components of spontaneous firing, contained 23% of the total stimulus-related variance, 86% of which was contained in one dimension (85% positive weights). Thus, overlap in the spaces encoding sensory and behavioral variables arises only because both can change the mean firing rate of the population: the precise patterns of increases and decreases about this change in mean were essentially orthogonal (Figure 4E).

To visualize how the V1 population integrated sensory and behavior-related activity, we examined the projection of this activity onto three orthogonal subspaces: a multidimensional subspace encoding only sensory information (stim-only); a multidimensional subspace encoding only behavioral information (behav-only); and the one-dimensional subspace encoding both (stim-behav shared dimension) (Figure 4F-G; Figure S10). During gray-screen periods there was no activity in the stim-only subspace, but when the stimuli appeared this subspace became very active. Conversely, activity in the behav-only subspace was present prior to stimulus presentation, and continued unchanged when the stimulus appeared. The one-dimensional shared subspace showed an intermediate pattern: activity in this subspace was weak prior to stimulus onset, and increased when stimuli were presented. Similar results were seen for the spont-only and stim-spont spaces (Figure 4F, lower panels). Across all experiments, variance in the stim-only subspace was 119 ± 81 SE times larger during stimulus presentation than during spontaneous epochs (Figure 4G), while activity in the shared subspace was 19 ± 12 SE times larger; activity in the face-only and spont-only subspaces was only modestly increased by sensory stimulation (1.4 ± 0.13 SE and 1.7 ± 0.2 SE times larger, respectively).

To visualize how stimuli affected activity in these subspaces, we plotted population responses to multiple repeats of two example stimuli (Figure 4H-K). When projected into the stim-only space, the resulting clouds were tightly defined with no overlap (Figure 4H), but in the behav-only space, responses to the two stimuli were directly superimposed (Figure 4I). Variability within the stimulus subspace consisted of changes in the length of the projected activity vectors between trials, resulting in narrowly elongated clouds of points, consistent with previous reports of multiplicative variability in stimulus responses [32–35]. A model in which stimulus responses are multiplied by a trial-dependent factor accurately captured the data, accounting for 89% ± 0.1% SE of the variance in the stimulus subspace (Figure 4J). Furthermore, the multiplicative gain on each trial could be predicted from facial motion energy (*r* = 0.61 ± 0.02 SE, cross-validated), and closely matched activity in the shared subspace (*r* = 0.73 ± 0.06 SE, cross-validated; Figure 4K). Although ongoing activity in the behav-only space and visual responses in the stim-only subspace added independently, we did not observe additive variability within the stim-only space itself: an “affine” model also including an additive term did not significantly increase explained variance over the multiplicative model (*p >* 0.05, Wilcoxon rank-sum test).

## Discussion

Ongoing population activity in visual cortex reliably encoded a latent signal of at least 100 linear dimensions, and possibly many more. The largest dimension correlated with arousal and modulated about half of the neurons positively and half negatively. At least 16 further dimensions were related to behaviors visible by facial videography, which were also encoded across the forebrain. The dimensions encoding motor variables overlapped with those encoding visual stimuli along only one dimension, which coherently increased or decreased the activity of the entire population. Activity in all other behavior-related dimensions continued unperturbed regardless of sensory stimulation. Trial-to-trial variability of sensory responses comprised additive ongoing activity in the behavior subspace, and arousal-dependent multiplicative modulation in the stimulus subspace, resolving apparently conflicting findings concerning the additive or multiplicative nature of cortical variability [20, 32–35].

Our data are consistent with previous reports describing low-dimensional correlates of locomotion and arousal in visual cortex [5, 7–14], but suggest these results were glimpses of a much larger set of behavioral and cognitive variables encoded by ongoing activity patterns. We found that 16 dimensions of facial motor activity can predict 31% of the reliable spontaneous variance. The remaining dimensions and variance might in part reflect motor activity not visible on the face or only decodable by more advanced methods [36–39], or they might reflect internal cognitive variables such as motivational drives.

The fact that ongoing and visually-evoked activity overlap in only one dimension at first appears to contradict previous reports showing similarity of sensory responses to ongoing activity [2–4]. We suggest three possible explanations for this apparent discrepancy. First, our experiments looked at a slower timescale than most previous studies, which binned the data at 100 ms [3], or even 2 ms [4]. Second, even a single dimension of common rate fluctuation is sufficient for some previously-applied statistical methods to report similar population activity [40]. Third, we found that encoding of non-sensory variables continued throughout stimulus presentation. Thus, similar firing patterns during stimulation and ongoing activity need not imply recapitulation of sensory events, just that cortex continues to encode nonsensory variables during sensory stimulation.

The brainwide representation of behavioral variables suggests that information encoded nearly anywhere in the forebrain is combined with behavioral state variables into a mixed representation. We found that these multidimensional signals are present both during ongoing activity and during passive viewing of a stimulus. Recent evidence indicates that they may also be present during a decision-making task [41]. What benefit could this ubiquitous mixing of sensory and motor information provide? The most appropriate behavior for an animal to perform at any moment depends on the combination of available sensory data, ongoing motor actions, and purely internal variables such as motivational drives. Integration of sensory inputs with motor actions must therefore occur somewhere in the nervous system. Our data indicate that it happens as early as primary sensory cortex. This is consistent with neuroanatomy: primary sensory cortex receives innervation both from neuromodulatory systems carrying state information, and from higher-order cortices which can encode fine-grained behavioral variables [6]. This and other examples of pervasive whole-brain connectivity [42–46] may coordinate the brain-wide encoding of behavioral variables we have reported here.

## Acknowledgements

We thank Michael Krumin for assistance with the two-photon microscopes, and Andy Peters for comments on the manuscript.

This research was funded by Wellcome Trust Investigator grants (095668, 095669, 108726, and 205093) and by a grant from the Simons Foundation (SCGB 325512). CS was funded by a four-year Gatsby Foundation PhD studentship. MC holds the GlaxoSmithKline / Fight for Sight Chair in Visual Neuroscience. CS and MP are now funded by HHMI Janelia. NS was supported by postdoctoral fellowships from the Human Frontier Sciences Program and the Marie Curie Action of the EU (656528).

## Author Contributions

Conceptualization, C.S., M.P., N.S., M.C. and K.D.H.; Methodology, C.S. and M.P.; Software, C.S. and M.P.; Investigation, C.S., M.P., N.S. and C.B.R.; Writing, C.S., M.P., N.S., M.C. and K.D.H; Resources, M.C. and K.D.H. Funding acquisition, M.C. and K.D.H.

## Supplementary Materials

### Materials and methods

All experimental procedures were conducted according to the UK Animals Scientific Procedures Act (1986). Experiments were performed at University College London under personal and project licenses released by the Home Office following appropriate ethics review.

All two-photon calcium imaging data is available at https://figshare.com/articles/Recordings_of_ten_thousand_neurons_in_visual_cortex_during_spontaneous_behaviors/6163622, in the form of processed deconvolved calcium traces together with the behavioral traces. All of the code used to analyze the data and produce the figures is available at www.github.com/MouseLand/stringer-pachitariu-et-al-2018a.

#### Preparation for two-photon calcium imaging in visual cortex

The imaging methods were similar to those described elsewhere [11]. Briefly, surgeries were performed in seven adult mice (P35–P125) in a stereotaxic frame and under isoflurane anesthesia (5% for induction, 0.5-1% during the surgery). We used mice bred to express GCaMP6s in excitatory neurons (1 EMX-CRE x Ai94 GCaMP6s mouse, 3 CamKII x tetO GCaMP6s mice, and 1 Rasgrf-CRE x Ai94 GCaMP6s mouse), or mice bred to express tdTomato in GAD+ inhibitory neurons (2 GAD-Cre x tdTomato mice), allowing inhibitory neurons to be identified and excluded from further analysis. We did not observe epileptiform activity in any of these mice [47].

Before surgery, Rimadyl was administered as a systemic analgesic and lidocaine was administered locally at the surgery site. During the surgery we implanted a head-plate for later head-fixation, made a craniotomy of 3-4 mm in diameter with a cranial window implant for optical access, and, in Gad-Cre x tdTomato transgenics, performed virus injections with a beveled micropipette using a Nanoject II injector (Drummond Scientific Company, Broomall, PA 1) attached to a stereotaxic micromanipulator. We used AAV2/1-hSyn-GCaMP6s, acquired from University of Pennsylvania Viral Vector Core. Injections of 50-200 nl virus (1-3 x1012 GC/ml) were targeted to monocular V1, 2.1-3.3 mm laterally and 3.5-4.0mm posteriorly from Bregma. To obtain large fields of view for imaging, we typically performed 4-8 injections at nearby locations, at multiple depths (~500 *μ*m and ~200 *μ*m). Rimadyl was then used as a post-operative analgesic for three days, delivered to the mice via their drinking water.

#### Data acquisition

We optically recorded neural activity in head-fixed awake mice implanted with 3-4 mm cranial windows centered over visual cortex, obtaining ~10,000 neurons in all recordings. The recordings were performed using multi-plane acquisition controlled by a resonance scanner, with planes spaced 30-35 *μ*m apart in depth. 10 or 12 planes were acquired simultaneously at a scan rate of 3 or 2.5 Hz. The mice were free to run on an air-floating ball and were surrounded by three computer monitors. Spontaneous activity was recorded in darkness (monitors off), or with a gray background or presented visual stimuli on these monitors arranged at 90*°* angles to the left, front and right of the animal, so that the animal’s head was approximately in the geometric center of the setup.

For each mouse imaged, we typically spent the first imaging day finding a suitable recording location, where the following three conditions held:

- the GCaMP signal was strong, in the sense that clear transients could be observed in large numbers of cells
- a large enough field of view could be obtained for 10,000 neuron recordings,
- the receptive fields of the neuropil were clearly spatially localized on our three monitors.

In animals for which there was a choice over multiple valid recording locations, we chose either: 1) a horizontally and vertically central retinotopic location or 2) a lateral retinotopic location, at 90*°* from the center, but still centered vertically. We did not observe differences related to retinotopic location (central or lateral), and thus pooled data across recording locations. We also did not observe significant differences between recordings obtained from GCaMP transgenic animals and from virus injections, nor between recordings made in complete darkness or with a gray screen. Thus, we pooled data over all conditions.

#### Visual stimuli

We presented 96 repetitions of 32 flashed natural images, covering all three screens. The images were manually selected from the ImageNet database ([48]), from ethologically-relevant categories: “birds”, “cat”, “flowers”, “hamster”, “holes”, “insects”, “mice”, “mushrooms”, “nests”, “pellets”, “snakes”, “wildcat”. We chose images if the subjects tended to fill out the image (less than 50% of the image was a uniform background), and if the images contained a balanced mixture of low an high spatial frequencies. The images were flashed for 0.5 sec with a randomized inter-stimulus interval between 0.3 sec and 1.1 sec, during which a gray screen was presented.

#### Calcium imaging processing

The pre-processing of all raw calcium movie data was done using a toolbox we developed called Suite2p, using the default settings [24]. The software is available at www.github.com/cortex-lab/Suite2P.

Briefly, Suite2p first aligns all frames of a calcium movie using 2D rigid registration based on regularized phase correlation, subpixel interpolation, and kriging. For all recordings we validated the inferred X and Y offset traces, to monitor any potential outlier frames that may have been incorrectly aligned. In a few recordings, a very small percentage (<0.01%) of frames that had registration artifacts were removed and the extracted traces were replaced with interpolated values at those frames. In all recordings, the registered movie appeared well-aligned by visual inspection. Next, Suite2p performs automated cell detection and neuropil correction. To detect cells, Suite2p computes a low-dimensional decomposition of the data, which is used to run a clustering algorithm that finds regions of interest (ROIs) based on the correlation of the pixels inside them. The extraction of ROIs stops when the pixel correlations of new potential ROIs drops below a threshold parameter, which is set as a fraction of the correlation in the high SNR ROIs; thus, it does not require the number of clusters to be set a priori. A further step in the Suite2p GUI classifies ROIs as somatic or not. This classifier learns from user input, reaching 95% performance on this data [24], thus allowing us to skip the manual step altogether for most recordings. We note that the 5% errors might be attributable to human labelling error, or to dendritic signals from backpropagating APs, reflecting the spiking of deeper cells. Thus, there is little risk of ROIs measuring signals other than neuronal action potentials.

We took great care to compensate cellular fluorescence traces for the surrounding neuropil signal [49]. This contamination is typically removed by subtracting out from the ROI signal a scaled-down version of the neuropil signal around the ROI; the scaling factor was set to 0.7 for all neurons. Importantly, for computing the neuropil signal, we excluded all pixels that Suite2p attributed to an ROI, whether somatic or dendritic. After neuropil subtraction, we subtracted a running baseline of the calcium traces with a sliding window of 60 seconds to remove long timescale drift in baseline, then applied non-negative spike deconvolution using the OASIS algorithm with a fixed timescale of calcium indicator decay of 2 seconds [50, 51]. To further ensure out-of-focus fluorescence could not contribute to our results, we excluded neurons whose signal might span two planes by excluding neurons in sequential planes that had a greater than a 0.6 correlation (in 1.2 second bins) with each other, and whose centers were within 5 *μ*m of each other in XY.

In addition, we ensured the cell sets used for reliable variance estimation (Figure 1H) were spatially non-overlapping: we segregated the field of view into 16 strips in XY (encompassing all Z) of width 60 *μ*m, and put cells from the odd strips in one group and the cells in the even strips in the other group. This ensured that no cells from different groups were at the same XY position but at a different depth. For peer prediction analyses (Figure S8), we excluded all peer cells within 70 *μ*m of the target (Euclidean distance in three-dimensional space).

#### Facial videography

Infrared LEDs (850nm) were pointed at the face of the mouse to enable infrared video acquisition in darkness. The videos were acquired at 30Hz using a camera with a zoom lens and an infrared filter (850nm, 50nm cutoff). The wavelength of 850nm was chosen to avoid the 970nm wavelength of the 2-photon laser, while remaining outside the visual detection range of the mice.

Running speed was not monitored videographically, but rather by optical mice placed orthogonally to the air floating ball on which the mouse stood.

#### Automated extraction of orofacial behaviors of mice

We developed a toolbox with a GUI for videographic processing of orofacial movements of mice. The software is termed FaceMap, and is available at www.github.com/MouseLand/FaceMap. The processing time taken by the software scales linearly with the number of frames, and runs 4x faster than real-time on 30 Hz videos.

##### Motion processing of regions of interest

To extract defined behavioral variables (e.g. pupil diameter, whisking), we used a graphical user interface which allows manual selection of face areas. The user can choose any region of the frame in which to compute the total absolute motion energy, the SVDs of the absolute motion energy or the SVDs of the raw frames.

The absolute motion energy at each time *T* is computed as the absolute value of the difference between consecutive frames, resulting in a matrix **M** of size *N*_pixels_ × *N*_timepoints_. The whisker signal used in the current study (Figure 1F) was defined to be the total motion energy summed over all pixels in a manually-defined region covering the whisker pad.

##### SVD computation for large matrices

To extract a high-dimensional representation of the facial signal, the toolbox applies singular value decomposition (SVD) to the raw movie, the motion energy movie, or both. The computation is identical in both cases.

The movie matrices are too large to decompose in their raw form. To compute their SVD, we first split the movie **M** into temporal segments **M**_*i*_ of length ~1 minute, and compute the SVD of each segment individually. Since the number of pixels is very large (> 1 million), we compute the SVD of of each movie segment by computing the top 200 eigenvectors **V**_*i*_ of its time by time covariance matrix. We then compute the spatial projections of the segment onto these components, **U**_*i*_ = **M**_*i*_**V**_*i*_. Each matrix **U**_*i*_ consists of the left singular vectors of **M**_*i*_, scaled by the singular values and is thus a 200-dimensional summary of the segment **M**_*i*_, related via an orthogonal projection. To estimate the SVD of the entire movie, we concatenate the **U**_*i*_ for all segments of the movie, and re-compute the SVD: [**U**_1_…**U**_*n*_] = **USV**^⊤^. The matrix **U** represents the spatial components of the full movie, and we project the the movie onto the top 1000 components of it, to obtain their temporal profiles: **W**_motion_ = **U**^⊤^**M**.

##### Pupil processing

To compute pupil area, the user first defines a region of interest using the FaceMap interface. The minimum value in this region is subtracted from all pixels for robustness across illumination changes. The darkest pixels in this region, identified by a user-selected threshold, correspond to the pupil. We estimate the pupil center as the center of mass of these dark pixels: 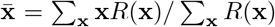, where **x** is the two-dimensional pixel location, *R*(**x**) is that pixel’s darkness level relative to the threshold, and the sum runs over all pixels **x** darker than the threshold. We compute the covariance of a 2D Gaussian fit to the region of interest: 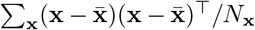, where the sum runs over all pixels darker than the threshold and *N*_x_ is the number of such pixels. For robustness, this process is iterated 4 times after re-selecting only pixels that are 2 standard deviations away from the center, and recomputing the Gaussian covariance fit. The final result is an outline of the pupil defined by an ellipse 2 standard deviations from the center of mass.

#### 8-probe Neuropixels recordings

Neuropixels electrode arrays [31] were used to record extracellularly from neurons in three mice. The mice were: 73 days old, male, and Drd1a-Cre(-/-) (mouse 1); 113 days old, female, and TetO-GCaMP6s;Camk2a-tTa (mouse 2); 99 days old, male, and Ai32;Pvalb-Cre (mouse 3). In all cases, a brief (<1 hour) surgery to implant a steel headplate and 3D-printed plastic recording chamber (~12mm diameter) was first performed. Following recovery, mice were acclimated to head-fixation in the recording setup. During head-fixation, mice were seated on a plastic apparatus with forepaws on a rotating rubber wheel. Three computer screens were positioned around the mouse at right angles. On the day of recording, mice were again briefly anesthetized with isoflurane while eight small craniotomies were made with a dental drill. After several hours of recovery, mice were head-fixed in the setup. Probes had a silver wire soldered onto the reference pad and shorted to ground; these reference wires were connected to a Ag/AgCl wire positioned on the skull. The craniotomies as well as the wire were covered with saline-based agar, which was covered with silicone oil to prevent drying. Each probes was mounted on a rod held by an electronically positionable micromanipulator (uMP-4, Sensapex Inc.) and was advanced through the agar and through the dura. Once electrodes punctured dura, they were advanced slowly (~10 *μ*m/sec) to their final depth (4 or 5 mm deep). Electrodes were allowed to settle for approximately 15 minutes before starting recording. Recordings were made in external reference mode with LFP gain=250 and AP gain=500, using SpikeGLX software. The mice were in a light-isolated enclosure and, during the part of the recording considered here, the computer screens were black. Data were preprocessed by re-referencing to the common median across all channels [52].

##### Spike sorting the Neuropixels data

We spike sorted the data using a modification of Kilosort [53], termed Kilosort2, that tracks drifting clusters. Code will be made publicly available at or before the time of publication. Without the modifications, the original Kilosort and similar algorithms can split clusters according to drift of the electrode, which would confound our behavioral-related analyses. Kilosort2 tracks neurons across drift levels and for longer periods of time (~1 hour in our case). To further mitigate the effect of drift we used a conservative threshold, excluding from further analysis units whose maximal firing rate was more than twice their minimal firing rates, after smoothing with a Gaussian filter of standard deviation 500 seconds. This excluded ~20% of the units on average. The final single units used were from several cortical areas (visual: 628, sensorimotor: 475, frontal: 664, retrosplenial: 161), hippocampal formation (1371), striatum (353), thalamus (2882), midbrain (885).

#### Correlations

Pairwise correlations were computed after binning activity at 1.2-1.3 s (3 or 4 frames respectively for 12 and 10 plane recordings; 1.2 s bins for Neuropixels recordings). To compute shuffled correlations (Figure 1C), we circularly shifted each neuron’s activity in time by a random number of bins (at least ±1000), and correlated all the shifted traces with all the original traces.

#### Arranging rasters by correlation

To visualize high-dimensional structure in raw data, raster plots were sorted vertically along a 1d continuum so that nearby neurons were most correlated. To do this, the binned activity of each neuron was first z-scored, and electrode data was high-pass filtered (100 s Gaussian kernel; this was not necessary for 2p data as traces had already been high-passed in preprocessing). Neurons were sorted using a generalization of scaled k-means clustering, where the clusters are ordered along a 1D axis to have similar means to their nearby clusters. Neurons were initially ordered based on their weights onto the first principal component of population activity, and divided into 30 equal-sized clusters along this ordering. On each iteration, we computed the mean activity of each cluster, smoothed it across clusters with a Gaussian window, then reassigned each neuron to the cluster whose smoothed activity it was most correlated with. This process was repeated for 75 iterations. The width of the Gaussian smoothing window was held at 3 clusters for the first 25 iterations, then annealed to 1 over the following 50 iterations. On the final pass, we upsampled the neurons’ correlations with each cluster by a factor of 100 via kriging interpolation with a smoothing constant of 1 cluster. This allowed us to determine sub-integer assignments of neurons to clusters, resulting in a continuous distribution of neurons along a 1D axis. The algorithm is available, implemented in Python and MATLAB at www.github.com/MouseLand/RasterMap. We ran the MATLAB version on the data here.

Although the electrode data was high-pass filtered to compute sorting, we display the original raw activity in Figure 3E.

#### Shared Variance Component Analysis

The SVCA method gives an asymptotically unbiased lower-bound estimate for the amount of a neural population’s variance reliably encoding a latent signal. A mathematical proof of this is given in the appendix; here, we describe how the algorithm was implemented for the current study.

We first split the population into two spatially segregated populations. To do so, we divided the XY plane into 16 non-overlapping strips of width 60 *μ*m, and assigned the neurons in the even strips to one group, and the neurons in the odd strips to the other group, regardless of the neuron’s depth. Thus, there did not exist neuron pairs in the two sets that had the same XY position but a different depth, avoiding a potential confound that a neuron could be predicted from its own out-of-focus fluorescence.

Neural population activity was binned at 1.2-1.3 s resolution (see above), and each neuron’s mean activity was subtracted from its firing trace. We divided the recording into training and test timepoints (alternating periods of 72 s each), thereby obtaining four neural activity matrices: **F**_train_, **F**_test_, **G**_train_, and **G**_test_ of size *N*_neurons_ × *N*_timepoints_, where **F** and **G** represent activity of the two cell sets. We compute the covariance matrix between the two cell sets on the training timepoints as

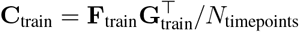

We then compute the top 1024 left and right singular vectors of **C**, yielding *N*_neurons_-dimensional vectors **u**_*k*_ and **v**_*k*_ for *k* = 1…1024. These vectors are the shared variance components (SVCs) for each population. The amount of reliable variance in each SVC (Figure 1L) is then estimated by the covariance of the SVC projections over the test samples:

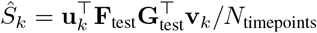

To obtain the fraction of reliable variance (Figure 1K), we normalize this reliable variance by the arithmetic mean of the variances of the test set data for each cell set on the corresponding projections, 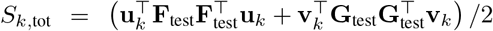.

#### Predicting neural activity from behavioral variables

To estimate the fraction of neural variance that could be predicted from explicitly-computed arousal variables (Figure 1M,N), we resampled their traces into the same 1.2-1.3 s bins as the neural data. The arousal variables (either single traces of running, whisking, pupil area or all three together) defined predictor matrices **X**_train_ and **X**_test_ for the training and test sets. We predicted the SVCs of neural activity **U**^⊤^**F**_train_ and **V**^⊤^**G**_train_ from the training-set behavior traces by unregularized multivariate linear regression, obtaining weight matrices **A** and **B** that minimized the squared errors ‖**U**^⊤^**F**_train_ − **AX**_train_‖^2^ and ‖**V**^⊤^**G**_train_ − **BX**_train_‖^2^. We then used these weight matrices to predict activity in the test set, and computed the covariance matrix of the residual error of each SVC:

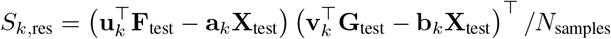

*S*_*k*,res_ represents the amount of variance along SVC *k* that cannot be predicted by the behavioral traces, and (*Ŝ*_*k*_ − *S*_*k*,res_)*/Ŝ*_*k*_ represents the fraction of reliable variance that can be so predicted. To compute the fraction of total variance explainable by behavioral traces (Figure 1M,N; Figure 2E,G-J), we normalize instead by the total test-set variance: (*Ŝ*_*k*_ − *S*_*k*,res_)*/S*_*k*,tot_.

To predict the fraction of neural variance that could be predicted from unsupervised videographic analysis (Figure 2), we took a similar approach, but computed the weight matrices **A** and **B** by reduced-rank regression. Reduced-rank regression is a form of regularized linear regression, with the prediction weights matrix restricted to a specific rank [54], reducing the number of parameters and making it more robust to overfitting. Figure 2E shows the fraction of total variance in successive dimensions that can be predicted by rank-16 prediction, while Figure 2G shows how the predicted fraction of variance in the first 128 dimensions depends on the rank of the predictor.

#### Peer prediction analysis

The shared variance component analysis described above – like a related algorithm for estimating reliable stimulus coding [29] – provides unbiased estimates, but requires thousands of simultaneously-recorded neurons per brain area. Because this many neurons were not available in our Neuropixels recordings, we turned to another method to estimate the reliable variance in these data. This method is an adaptation of the previously-described “peer prediction” method [55, 56]. Peer prediction analysis attempts to predict each neuron individually from the other simultaneously recorded cells (the neuron’s “peers”). In contrast, SVCA finds the dimensions of activity in a large population that can be most reliably predicted from a held-out set of neurons. Because a substantial fraction of a single neuron’s variance arises from independent noise, which is averaged out when projecting onto the SVCA dimensions, peer prediction gives systematically lower values of variance explained than SVCA.

To apply peer prediction to our data, we again binned neural activity with 1.2-1.3 s resolution, and divided these timepoints into a training set and a test set, consisting of alternating blocks of duration 72 s. Each neuron took a turn as target for prediction from the activity of simultaneously recorded “peer” cells, defined to be any cells on all other probes and cells on the same probe greater than 5 sites away (40 *μ*m) for neuropixels recordings; for 2p recordings we used all neurons greater than 70 *μ*m from the cell in 3D distance, in order to avoid potential optical contamination from the target neuron. We denote peer cell activity in the training and test sets by *N*_cells_ × *N*_timepoints_ matrices **F**_train_ and **F**_test_, respectively, and target cell activity in the training and sets as 1 × *N*_timepoints_ vectors **g**_train_ and **g**_test_. We first computed the singular value decomposition of peer cell activity on the training set: **F**_train_ = **USV**^⊤^. We then predicted the target neuron activity by ridge regression from *n* singular value components of peer cell activity, where *n* took values *n* = 1, 2, 4, 8, 16, …, 512, 1024. The prediction weights were thus

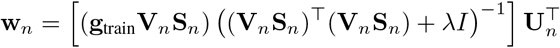

where **U**_*n*_, **V**_*n*_, **S**_*n*_ are matrices containing the top *n* singular vectors. Then the prediction of the single neuron activity on the test set was 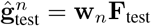, and the fraction of variance explained was 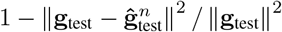. We chose to be 10 by hand.

#### Subspaces of stimulus and behavioral activity

The stimulus subspace (Figure 4) was defined as the space spanned by the trial-averaged responses of each of the 32 stimuli presented. We computed the *N*_neurons_ × *N*_stimuli_ matrix of trial-averaged responses **R** from one third of the stimulus responses, saving the other two thirds of the stimulus responses for variance estimation.

The behavior subspace was defined via the reduced-rank regression prediction method described above. We performed the regression on one-half of the spontaneous activity, leaving the other half of the spontaneous activity for variance estimation. This method produces a weight matrix of size *N*_neurons_ × *N*_facePCs_, that factorizes as a product of two matrices of sizes *N*_neurons_ × *r* and *r × N*_facePCs_, where *r* is the rank of prediction. We defined the behavior space as the space spanned by the first 32 columns of the former matrix, and we define a *N*_neurons_ × 32 matrix **E**_*B*_ whose columns contain an orthonormal basis for this space.

To determine dimensions inside the behavioral subspace that contain stimulus information, we found the sequence of orthogonal directions **e**_*i*_ maximizing the sum of squared projections (the power) of the trial-averaged stimulus responses **R**:

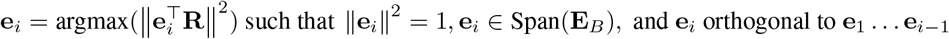

The solution to this maximization problem is given by the left singular vectors of **E**^⊤^_*B*_**R**. To determine the amount of stimulus variance in shared dimension *i* (Figure 4C), we projected test data onto **e**_*i*_ and quantified the stimulus-related variance in this projection using the unbiased method of [29] on the remaining two thirds of stimulus responses. We call **e**_1_ the “shared stim-behav” dimension, because it contains significant stimulus variance, as opposed to **e**_2_, **e**_3_, … which contain very little (Figure 4C). A histogram of the weights of **e**_1_ on all neurons are plotted in Figure 4D.

The behav-only subspace was defined by projecting out the shared dimension **e**_1_ from all columns of **E**_*B*_. The stim-only subspace was defined by projecting out from all rows of **R** the top right singular vector of the matrix **E**^⊤^_*B*_**R**. Timecourses of neural activity projected into these subspaces is plotted in Figure 4F. To quantify the amount of variance in each subspace (stim-only, behav-only, stim-behav) (Figure 4G), computed the total projected variance as the sum of squared projection lengths along each axis of an orthonormal basis.

An identical analysis was used to define the stimulus-spontaneous shared dimension, and spontaneous-only subspace, by replacing the subspace of the top 32 behavioral components with the subspace of the top 128 principal components of activity computed in one half of the spontaneous period.

To account for trial-to-trial variability in the stimulus subspace, we fit a multiplicative gain model. A gain parameter *g*_*t*_ was fit for each trial *t*, and the activity of dimension *n* in response to the stimulus *σ*_*t*_ shown on this trial was modelled as 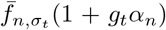. Here, 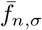 represents the mean activity of dimension *n* to stimulus *σ*, and *α*_*n*_ represents the susceptibility of this dimension to gain fluctuations. Note that the mean of *g*_*t*_ across trials from the same stimulus is 0 by definition. We used an alternating optimization method to obtain the best fit **g** given *α*, then *α* given **g**, repeating for 100 iterations. We also evaluated an affine model, allowing both the gain and offset of each neuron’s responses to change on a trial by trial basis: 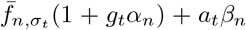. Here, *a*_*t*_ is an additive offset on trial *t*, with each neuron scaling this offset by a factor *β*_*n*_. The vector *β* can therefore describe directions of additive variability inside the stimulus subspace.

### Mathematical Appendix

#### Proof that SVCA is an asymptotically unbiased estimate of reliable variance

Here we describe the SVCA method in more detail, and prove that the reliable variance it estimates has an expectation that is a lower bound for the true value, with bias vanishing in the limit of many neurons and stimuli.

The key to the SVCA method is to split spontaneous neuronal activity into a reliable component – which coherently and deterministically encodes an unobservable state variable – and an unreliable component, which is random and independent between neurons. The unreliable component is sometimes referred to as “Poisson noise”, as it would occur in a model where neurons fired a number of spikes drawn from a Poisson distribution whose rate deterministically encoded the state variable. Such behavior can also occur in networks where strong and balanced excitation and inhibition lead to chaotic variations in spike counts even in deterministic simulations [12, 21, 57].

As we show in the main text, spontaneous activity encodes details of ongoing facial behavior – but the reliable component of spontaneous activity can also represent internal cognitive variables that cannot be directly measured. SVCA allows one to estimate the amount of neuronal population variance that coherently and reliably encodes an unobservable state variable, distinguishing it from the variance that is independently variable between neurons.

Formally, we consider the reliable component of spontaneous population activity to represent a state variable *s*, which is drawn from a set of possible states 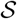 according to a probability distribution 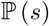. We model the activity of neuron *n* at timepoint *t* as the sum of a reliable response 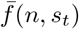 that depends deterministically on the state *s*_*t*_ at this timepoint, plus a noise term *ϵ*_*n,t*_ which is independent between neurons, and independent of the state *s*_*t*_:

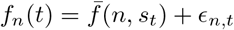

To estimate the variance of reliable population activity, we divide the recorded neurons into two sets of size *N* and *M*. We denote the responses of the first set on trial *t* by *f*_*n*_(*t*) for *n* = 1 … *N*, and the second set by *g*_*m*_(*t*) with *m* = 1 … *M*, summarizing their activity in random vectors 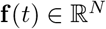 and 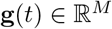. Define the expected *N × M* cross-covariance matrix of these vectors as **C**, with entries 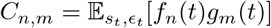 which is equal to 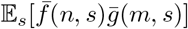 by independence of the noise terms. We write its singular value decomposition as 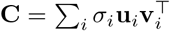, where the *σ*_*i*_ are non-increasing real numbers, and the **u**_*i*_ and **v**_*i*_ are orthonormal systems in 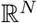 and 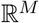 respectively.

To estimate the amount of reliable variance in an optimal *k*-dimensional subspace, we compute the first *k* left and right singular vectors **û**_*i*_ and 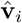 from the covariance matrix estimated from training timepoints, and estimate the *k*-dimensional reliable variance as

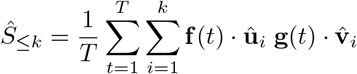

where the first sum runs over test set trials *t*. Because the training and test sets are independent, and because *ϵ* is independent of *s*,

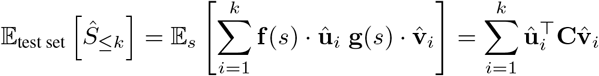

Now, because**û**_*i*_ and 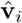 are orthonormal systems, 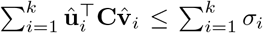, with equality when **û**_*i*_ and 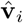 are the first *k* singular vectors of **C**, which they approach when the number of training timepoints becomes large. Thus, the expectation of *Ŝ*_≤*k*_ is a lower bound for 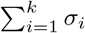, that becomes accurate in the limit of a large number of training timepoints.

Finally, we show that as the number of recorded neurons increases, 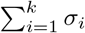 converges to the reliable variance of the entire population’s top *k* dimensions. Recall that the singular values of **C** are square roots of the eigenvalues of **CC**^⊤^. Concatenating the reliable rate vectors into *T × N* and *T × M* matrices **F** and **G** so **C** = **F**^⊤^**G**, we see that 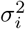 is the *i*^*th*^ eigenvalue of **F**^⊤^**GG**^⊤^**F**, which is also the *i*^*th*^ eigenvalue of **GG**^⊤^**FF**^⊤^ by cyclic permutability of eigenvalues. No matter how many neurons we record from, **FF**^⊤^ and **GG**^⊤^ are both *T × T* matrices, and we next consider their limit as the number of neurons becomes large.

To do so, we consider the neurons to be drawn from a hypothetically infinite population of neurons 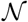 according to a probability distribution 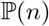. As the number *N* of neurons sampled from this distribution becomes large, the (*t*_1_, *t*_2_)^*th*^ entry of the matrix 1 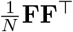 converges to

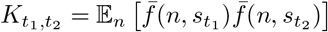

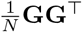 converges to the same limit as the number of neurons *M* in the second set becomes large. Thus, the *k* first singular values of **C** tend to the *k* first eigenvalues of **K**, whose sum is the amount of reliable variance in the optimal *k*-dimensional subspace of population activity.

---

**Figure S1.**
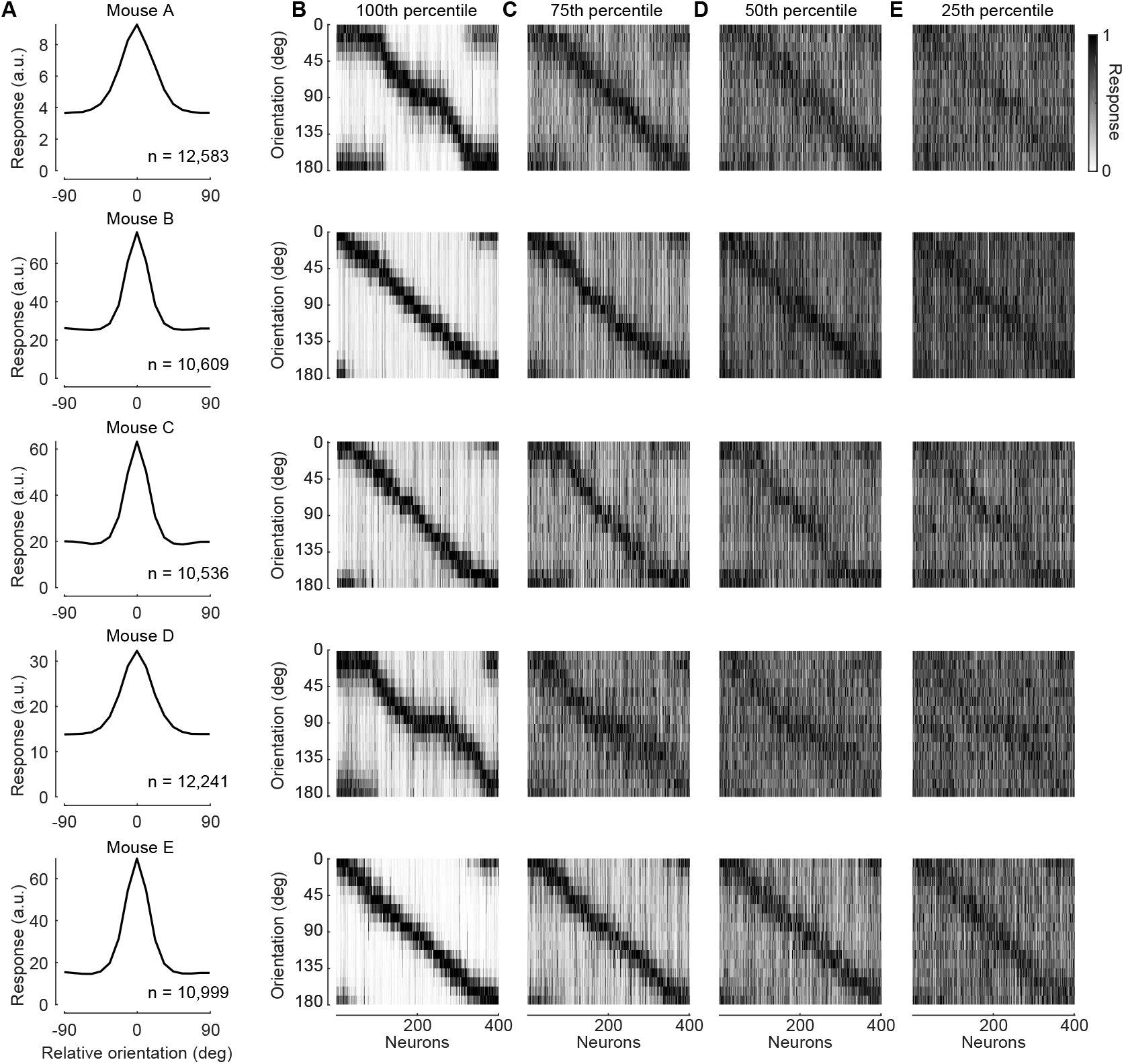
Robust orientation tuning in recorded V1 neurons. (A) Orientation tuning curve, showing population-averaged responses as a function of angle relative to each cell’s preferred orientation (obtained from held-out responses). Stimuli were Gabor drifting gratings of spatial and temporal frequency of .05 cpd and 2 Hz. Each row shows responses a single recording session from a single mouse. (B) Orientation tuning curves of the 400 most tuned neurons in each experiment (as assessed by orientation selectivity index), arranged horizontally by preferred orientation (computed from held-out responses). (C-E) Orientation tuning curves of 400 neurons taken from the 75th, 50th, and 25th percentile of orientation selectivity, displayed similarly.

**Figure S2.**
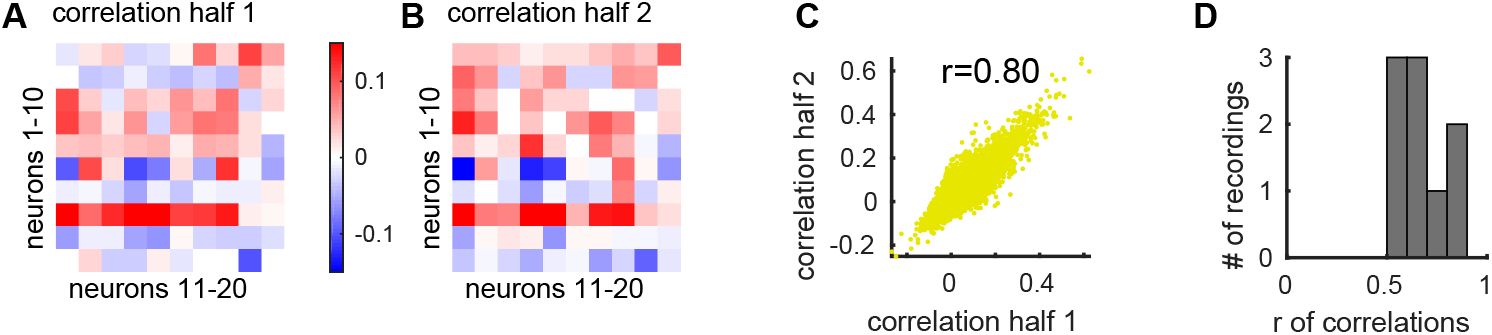
Correlation matrices are structured and consistent across time. (A,B) Pseudocolor representation of spontaneous correlation matrices for a subset of cells, computed independently from two halves of recording. (C) Scatter plot showing correlations of each cell pair in a single recording, for two independent halves. (D) Histogram showing Pearson correlation coefficient of of pairwise correlations (as in C), for all recordings.

**Figure S3.**
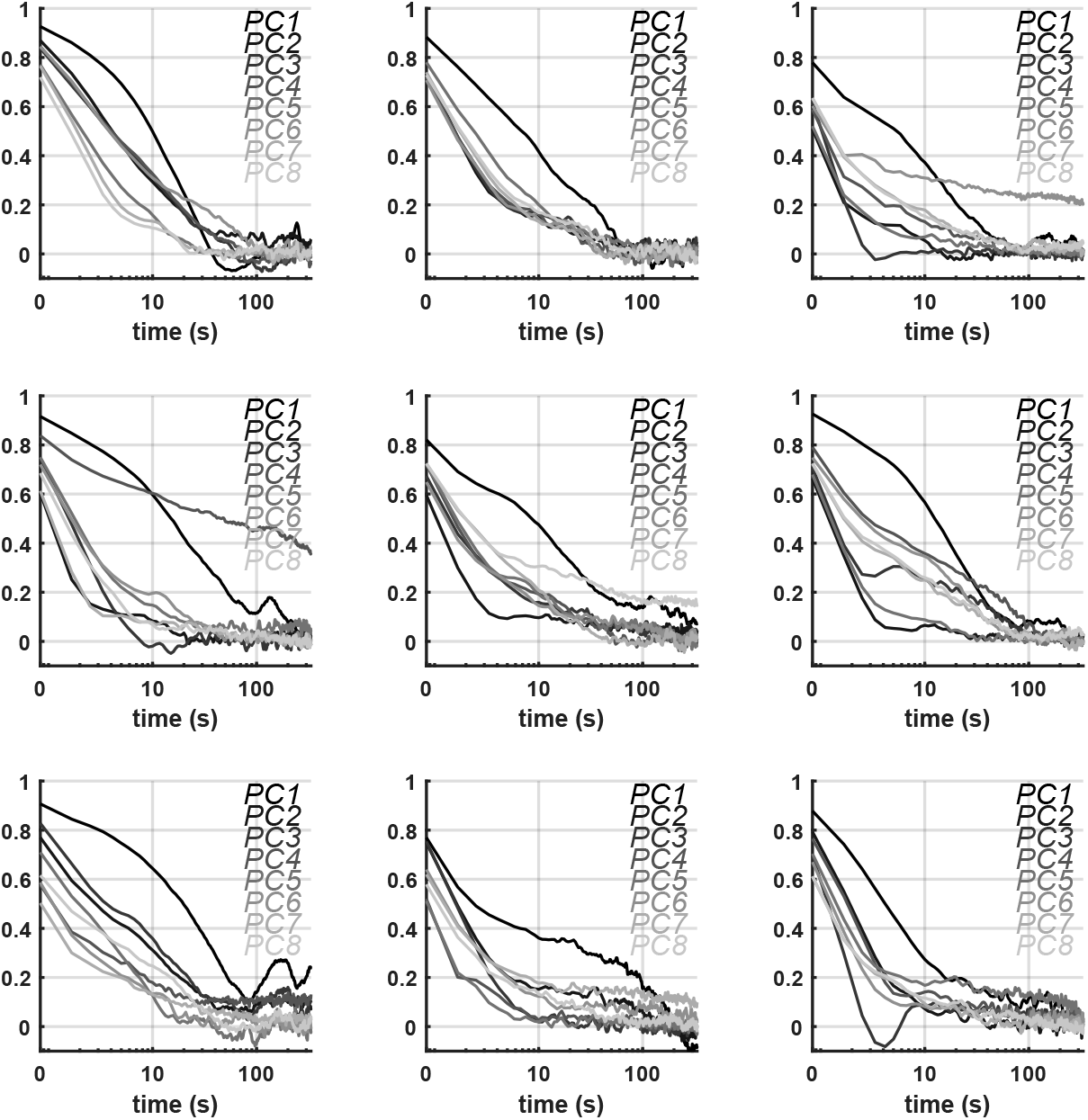
Temporal autocorrelation of ongoing neural activity. Each panel shows data for one recording; within each plot, each curve shows the temporal autocorrelation of a single principal component of ongoing population activity (1.2 second bins; log x-scale). Second plot on top row is for the example recording shown in Figure 1.

**Figure S4.**
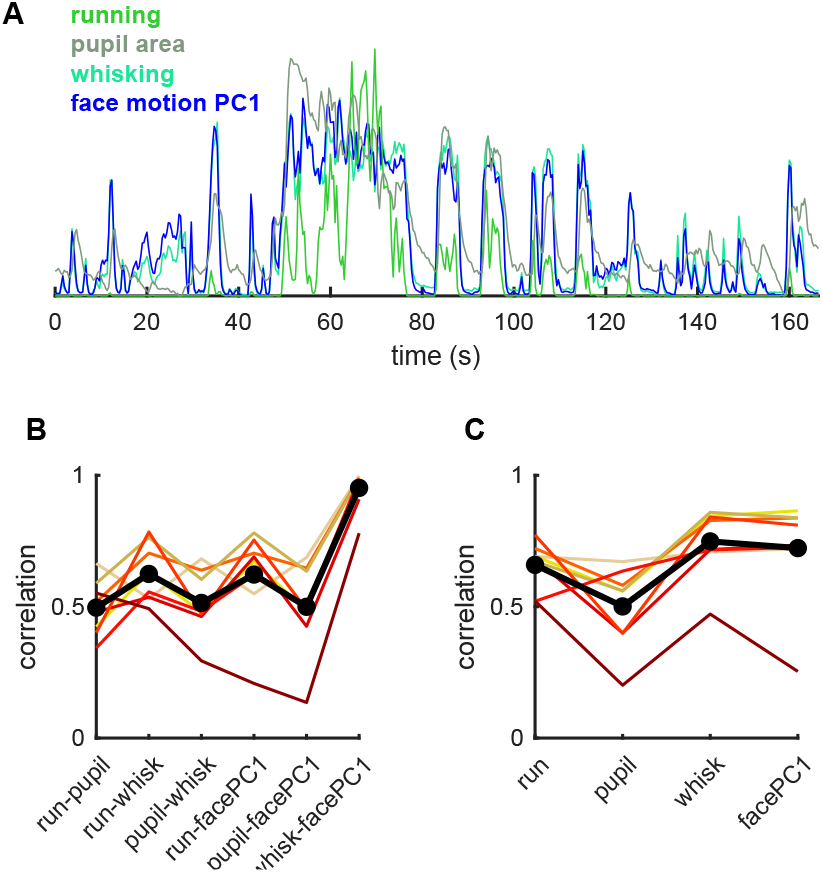
Correlations between arousal variables. (A) Time courses of running speed, pupil area, whisker motion energy, and first PC of motion energy, for a 160 s segment of an example experiment scaled to have the same maximum and minimum. (B) Pearson correlations for each pair of these variables. Each line denotes an individual experiment, black denotes average. (C) Correlation of the first neural PC with each behavioral variable.

**Figure S5.**
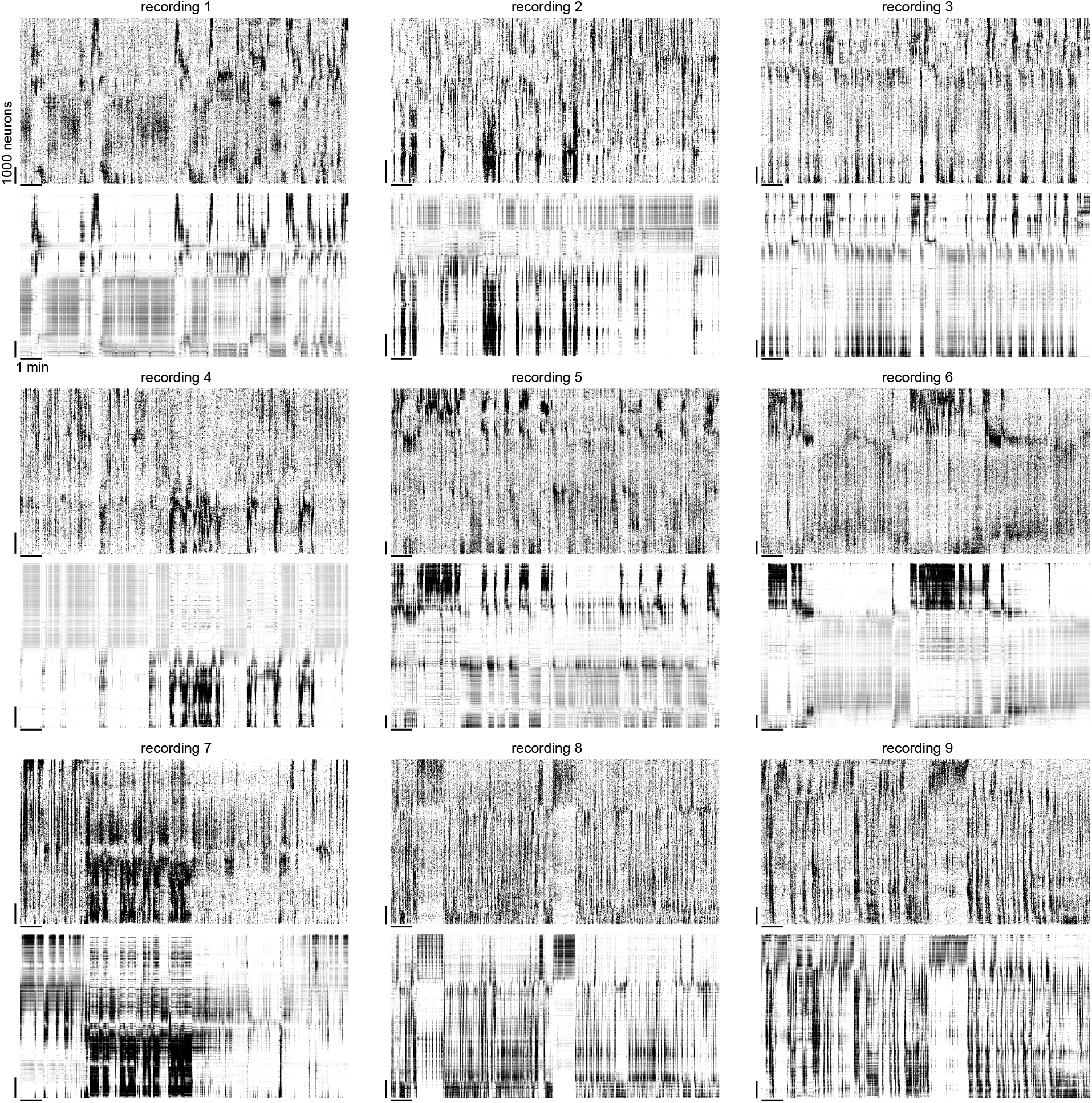
Additional examples of predicting high-dimensional neural population activity from facial videography. For each panel, the top plot shows a raster diagrams sorted vertically to place correlated neurons together (cf. Figure 1G). Bottom plots show predictions from facial videography (cf. Figure 2F).

**Figure S6.**
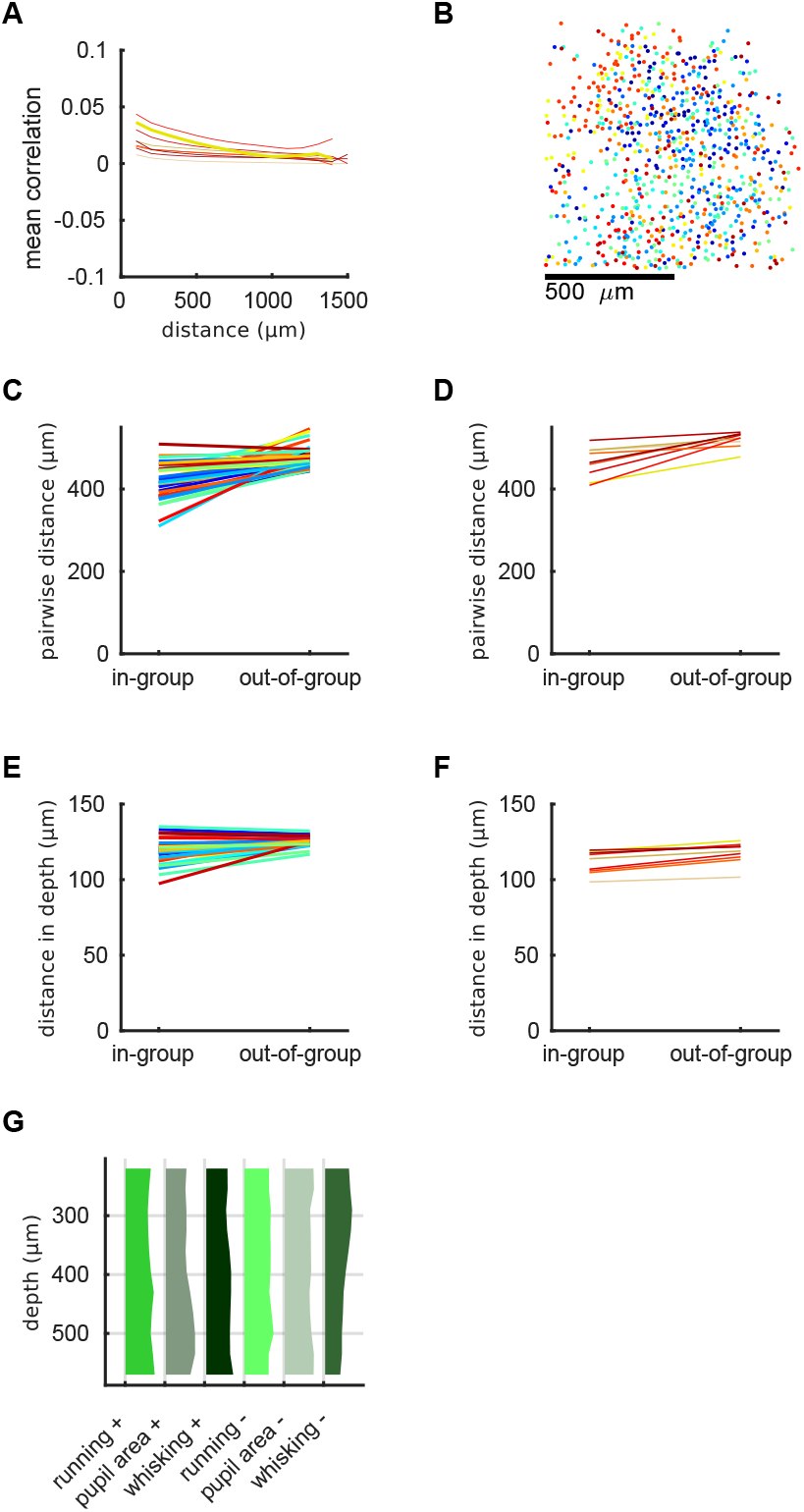
Spontaneous correlations bear little relationship to cortical distance. (A) Mean correlation over cell pairs, as a function of distance on the cortical surface. Each line represents a single experiment. (B) Neurons were assigned to 30 clusters according to a 1d manifold-embedding algorithm (Figure 1G). Points show XY locations of cells in one example imaging plane, colored according to cluster. (C) Three-dimensional distances between neurons in the same or different clusters, for one experiment. Each line represents a cluster. (D) Average of (C) over all clusters. Each line represents an experiment. (E,F) Same as (C,D) but for one-dimensional pairwise distances in depth. (G) Distribution across cortical depth of six non-overlapping groups of neurons with different behavioral correlates. Neurons were grouped according to which arousal variable they best correlated with and the sign of this correlation.

**Figure S7.**
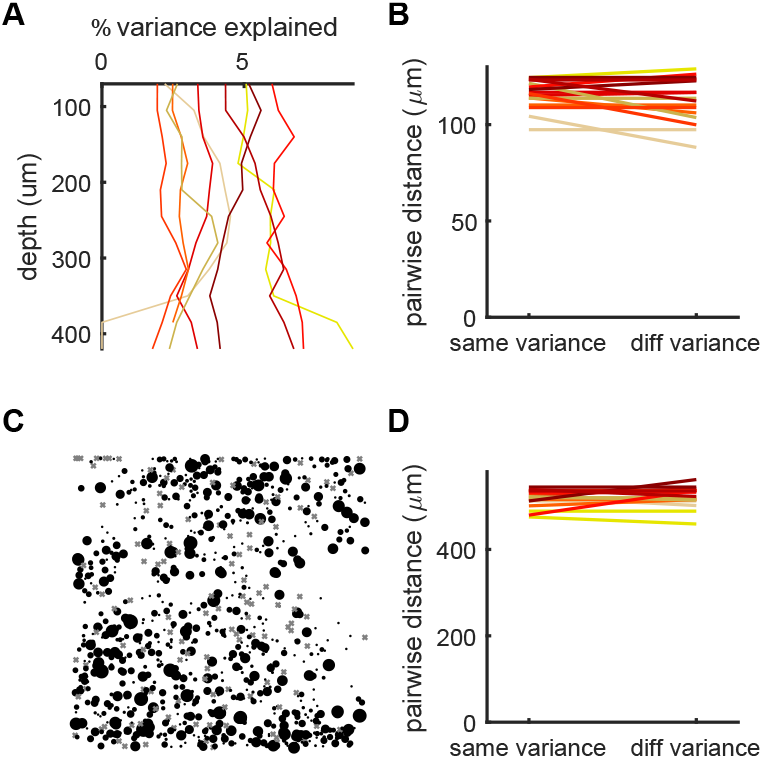
Predictability of neural activity from facial motion is spatially uniform in V1 recordings. (A) Percentage of single-neuron variance explained by facial motion, averaged over cells as a function of cortical depth (16-dimensional reduced rank regression, cross-validated, 1.2 second bins). Each line represents a different experiment. Note that single-neuron prediction explains a lower percentage of variance than SVC prediction (Figure S8). (B) Neurons were split into two groups: low variance explained (<3%) and high variance explained (>10%). The average vertical distance between neurons in the same group (“same variance”) was similar to the distance between neurons with different variance levels (“diff variance”) (115 *μ*m vs 117 *μ*m, *p >* 0.05 Wilcoxon rank-sum test). The lack of difference indicates that predictability from facial motion does not depend systematically on cortical depth. (C) Fraction of variance explained as a function of XY position in V1, for an example recording plane. Each dot represents a cell, of size proportional to the explained variance; crosses indicate cells of negative explained variance on the test set. (D) Same as B, but for XYZ distance. The lack of difference (516 *μ*m vs 525 *μ*m, *p >* 0.05 Wilcoxon rank-sum test) indicates that variance explained does not depend systematically on XYZ position.

**Figure S8.**
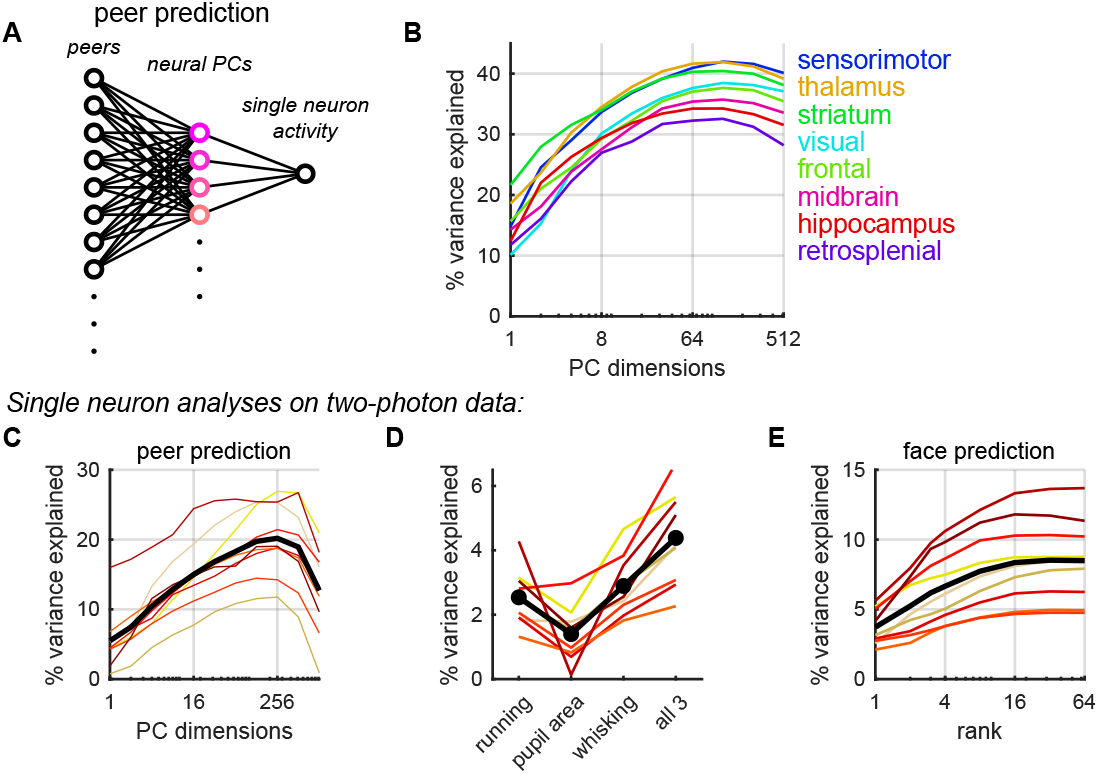
Explaining single neuron variance using peer prediction and behavioral variables. Shared variance component analysis – like a related algorithm for estimating reliable stimulus coding [29] – requires thousands of simultaneously-recorded neurons per brain area. Because this many neurons were not available in our Neuropixels recordings, we instead estimated the reliable variance predictable from facial behavior using an adaptation of the “peer prediction” method [55, 56]. (A) In this method, each neuron takes a turn as target cell, and principal component analysis is applied to the activity of its simultaneously recorded “peers” by ridge regression, excluding spatially neighboring neurons. The activity of the target cell is predicted from these principal components, and prediction quality is assessed via cross-validation. (B) Average single neuron variance explained by peer prediction as a function of the number of principal components, for each brain area (neurons pooled across 3 mice). (C) Similar analysis for two-photon calcium imaging in V1. The peak predictability of 20.2% ± 1.7% of variance explained is obtained when predicting from 256 peer PCs; because the independent noise in a single neuron’s activity cannot be predicted from other neurons, this is substantially lower than the 97% reliability of the first SVC, and the 67% reliability of the first 128 SVCs together. (D) Predicting single neuron activity from arousal variables; because the independent noise in a single neuron’s activity cannot be predicted from behavior, predictability is lower than when predicting SVCs (cf. Figure 1N). (E) Predicting single neuron activity from multidimensional behavior information using reduced rank regression (cf. Figure 2G).

**Figure S9.**
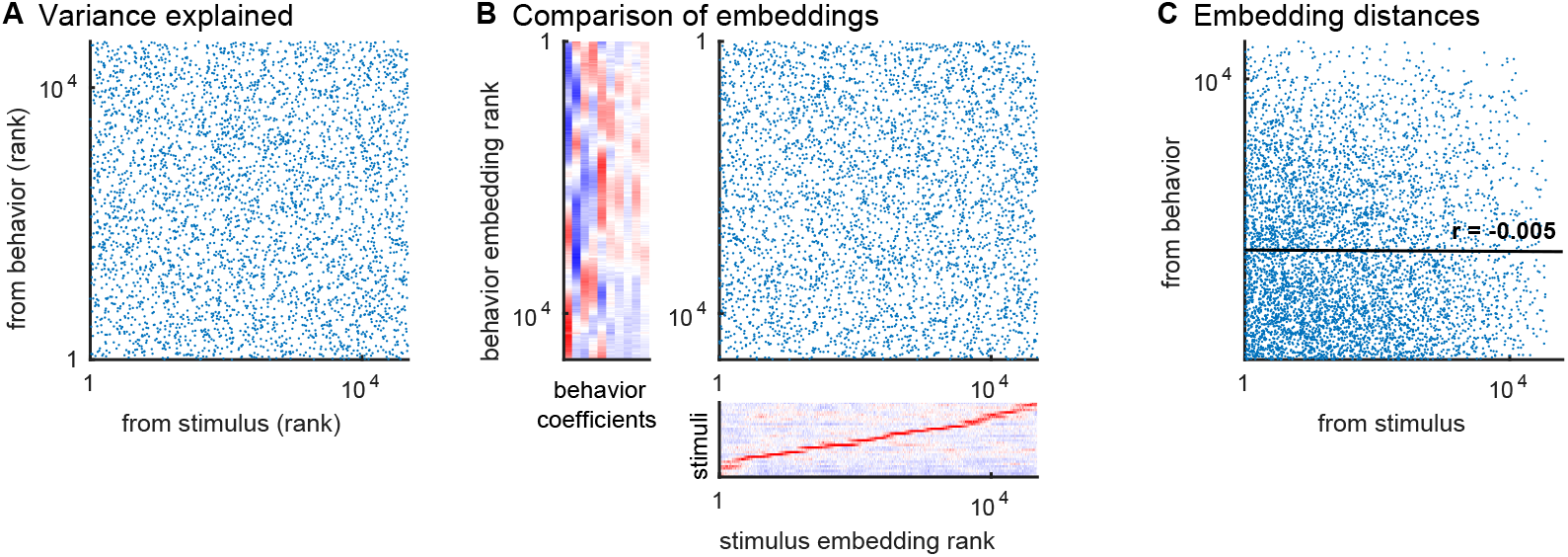
Mixed representation of stimuli and behavior in visual cortex. (A) Comparison of variance explained by stimuli and by behavior videography for single neurons. X- and y-coordinates represent each neuron’s rank order from least to most predictable by stimulus and behavior, respectively. The extremely weak correlation (r = -0.18, p<0.01 Spearman’s rank correlation) indicates that the strength with which a neuron is modulated by stimuli is essentially unrelated to the strength with which it is modulated by behavior. (B) Comparison of tuning to behavior and to stimuli. Each point represents a neuron, and the x- and y-coordinates represent its rank in two separate 1D embeddings computed from stimulus responses, and from behavioral coefficients of the face prediction model (16D), respectively. (C) Pairwise comparison of visual-tuning similarity (x-axis) and behavioral-tuning similarity (y-axis). Each point represents a pair of cells; tuning similarities are defined as the pairwise distance in the 1D embeddings on the x- and y-axes of panel B. The lack of correlation (r = -0.005, p > 0.05) indicates that neurons which are tuned for similar stimuli are no more likely to be tuned for similar behavioral features.

**Figure S10.**
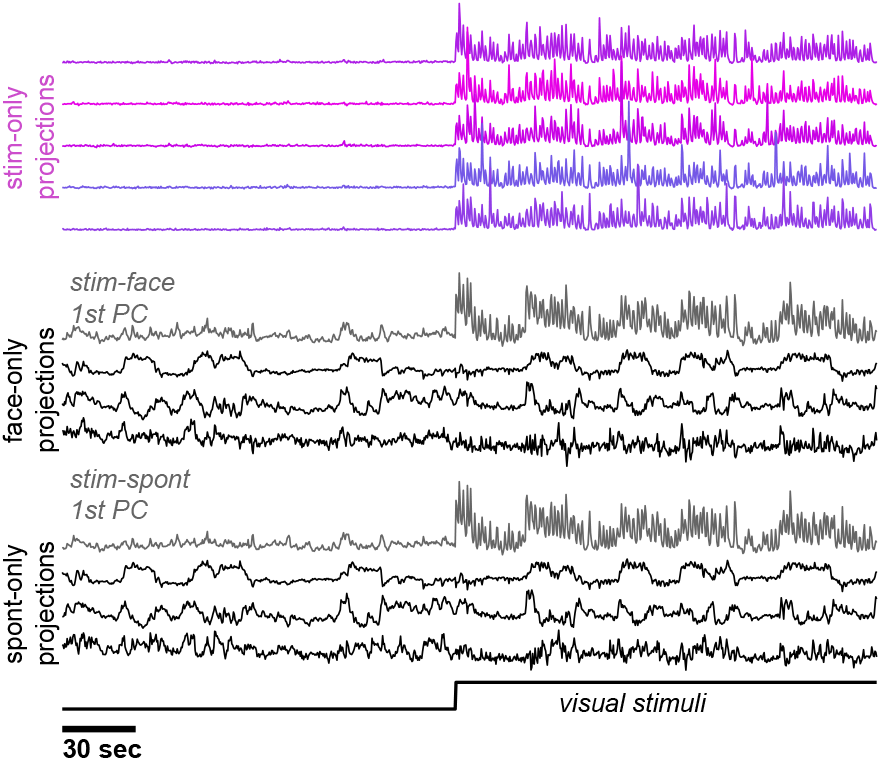
Short-timescale view of projections onto activity subspaces, corresponding to a zoom into Figure 4F.

**Movie S1. Spontaneous neural activity of 10,000+ neurons in visual cortex of awake mice.** Two-photon calcium imaging of 11 planes spaced 35 *μ*m apart. Movie speed is 10x real-time.

**Movie S2. Multi-dimensional spontaneous behaviors.** Movie speed is 5x real-time.

**Movie S3. Spontaneous behaviors are correlated with spontaneous neural activity.** Video of mouse face recorded simultaneously with neural activity. Movie speed is 10x real-time.

